# A PthXo2B ortholog in *Xanthomonas oryzae* pv oryzae strain IX-221 acts as a major virulence factor on *indica* rice without activating a Clade III *SWEET* gene

**DOI:** 10.1101/2024.08.08.607113

**Authors:** Prashant Mishra, Shakespear S, Sara C D Carpenter, Hamsa S, Vigi S, Anith K N, Prasanta K Dash, Adam J Bogdanove, Rhitu Rai

**Affiliations:** Plant Pathogen Interaction, ICAR-National Institute of Plant Biotechnology, Pusa, New Delhi, India; Plant Pathology and Plant-Microbe Biology section, School of Integrative Plant Science, Cornell University, Ithaca, New York, USA; Department of Microbiology, College of Agriculture, Kerala Agricultural University, Vellayani, Thiruvanathpuram, Kerala, India

**Keywords:** *Xanthomonas oryzae*, transcription activator-like effector (TALE), aberrant repeat, *xa13*, bacterial blight of rice, susceptibility gene, SWEET sugar transporter

## Abstract

To infect rice, *Xanthomonas oryzae* pv. oryzae (*Xoo*) deploys transcription <u>a</u>ctivator-like <u>e</u>ffectors (TALEs) that specifically bind and upregulate host “susceptibility” (S) genes. 34-amino acid (aa) repeats in TALEs interact one-to-one with DNA bases. Variation at positions 12 and 13 in each repeat, the repeat-variable diresidue (RVD), determine specificity. Some repeat variants shorter or longer than 34 aa can disengage to accommodate a single base deletion in the target sequence. *OsSWEET11*, *13*, and *14* are key S genes, targeted by different TALEs from diverse *Xoo* strains. *xa13* is a *SWEET11* allele lacking the TALE binding site and thus conferring resistance. *xa13* is overcome by TALEs that activate *SWEET13* or *SWEET14*. We report here that an *xa13*-compatible *Xoo* strain, IX-221, from India, harbours an ortholog of the *SWEET14*-targeting TALE PthXo3 and two orthologs of the *SWEET13*-cognate PthXo2, each with one or two 36-aa repeats capable of disengaging. One of the PthXo2 orthologs, PthXo2B_IX-221_, has a repeat region identical to the previously characterized PthXo2B_PXO61_, except for a two amino acid difference near the end of the 19^th^ repeat. Like PthXo2B_PXO61_, PthXo2B_IX-221_ upregulates *SWEET13* in japonica rice and no *SWEET* in *indica* rice, but unlike PthXo2B_PXO61_ it nonetheless renders *indica* rice susceptible, pointing to an alternative *S* gene. Further, a designer TALE (dTALE) constructed using a standard, consensus sequence for each repeat and RVDs identical to those of PthXo2B_IX-221_ failed to render *indica* rice susceptible. Alignment of the PthXo2B_IX-221_ repeats shows a departure from the consensus in each of two repeats carrying the RVD ‘NN’: the sequence ‘MAIAN’ in place of ‘VAIAS’ beginning at position 7. Together, the PthXo2B_IX-221_ results thus suggest that non-RVD sequence variation affects TALE targeting profiles. More broadly, the presence of the three aberrant repeat-harbouring TALEs in IX-221 suggests that widespread deployment of *xa13* in India resulted in strains super-equipped to overcome it, capable of activating multiple *SWEET* genes and alleles as well as an apparent alternate S gene.

## Introduction

*Xanthomonas oryzae* pv. oryzae (*Xoo*) causes bacterial blight disease of rice (BB). It is important economically and as a model to study host-pathogen interaction and coevolution (Hutin *et al*., 2015, Nino-Liu *et al*., 2006). *Xoo* injects DNA-binding proteins called transcription activator-like effectors (TALEs) into rice cells, where they localize to the nucleus and increase the transcription of host genes by binding to effector-specific promoter sequences called effector binding elements (EBE) (Bogdanove *et al*., 2010). Those genes that contribute to disease development when so upregulated are considered susceptibility (S) genes. Binding of an EBE by a TALE is governed by a central repeat region (CRR) of the protein composed of nearly identical direct repeats of 33-35 amino acids (aa), differing at the 12^th^ and 13^th^ positions, together called the repeat variable diresidue (RVD). Following a partially degenerate code, each RVD directly interacts with a single nucleotide, such that the number and composition of RVDs predicts the sequence of the EBE (Moscou & Bogdanove, 2009, Boch *et al*., 2009). Besides the standard 33-35 aa repeats, variants with repeat lengths of 28, 30, 36, 39, 40 and 42 aa have been reported (Richter *et al*., 2014b, Wilkins *et al*., 2015, Oliva *et al*., 2019b). Some of these so-called aberrant repeats can function as a standard repeat or disengage in an as yet structurally uncharacterized way to accommodate a single base pair deletion at the corresponding location in the target (Richter et al., 2014b, Becker *et al*., 2022).

A major class of S gene targeted by TALEs in BB consists of clade III members of the ‘SWEET’ sucrose transporter gene family (Streubel *et al*., 2013). The cognate TALEs are referred to as major TALEs, owing to their important contribution to virulence as activators of these major S genes. The first such TALE identified was PthXo1, which induces *SWEET11*, also called *Os8N3* or *Xa13* (Chu *et al*., 2006, Yang *et al*., 2006). Recessive alleles of *SWEET11*, collectively known as *xa13*, can confer resistance (rather, loss of susceptibility) to strains that depend on PthXo1, by virtue of any of several promoter mutations that disrupt the PthXo1 binding site. These *xa13 SWEET11* alleles are ineffective against *Xoo* strains with TALEs that activate other Clade III *SWEET* genes, such as PthXo2, which activates *SWEET13* (Zhou *et al*., 2015, Oliva et al., 2019b), or AvrXa7, PthXo3, TalC, and TALE5, which activate *SWEET14* (Antony *et al*., 2010, Yu *et al*., 2011, Streubel et al., 2013). While PthXo2 activates *SWEET13* and acts as a major TALE only on *indica* rice, a variant named PthXo2B with aberrant 9^th^ and 12^th^ repeats of 36 aa, found in a few *Xoo* strains, activates the *japonica*, and not the *indica* allele of SWEET13 (Oliva et al., 2019b) and confers virulence only toward *japonica* rice. The key roles of *SWEET11*, *13*, and *14* in bacterial blight of rice, inspired the development of lines of select *indica* and *japonica* mega rice varieties edited at EBEs in all three to provide broad spectrum bacterial blight disease resistance (Oliva et al., 2019b, Xu *et al*., 2019).

The naturally occurring *SWEET11* allele *xa13* has been widely deployed in India, but the Indian *Xoo* population is quite diverse, and strains that overcome *xa13* have been reported from different parts of the country (Lore *et al*., 2011, Mishra *et al*., 2013, Yugander *et al*., 2017). (Midha *et al*., 2017, Mondal *et al*., 2014). In individual *Xoo* strains, TALEs are typically numerous (15 or more), and within and across strains they are diverse. Their abundance and the repetitive nature of their coding sequences likely contribute to the diversity, facilitating recombination and rapid adaptation under selection pressure (Booher *et al*., 2015, Denancé *et al*., 2018). Detailed molecular characterization of resistance-breaking strains is essential for insight into pathogen-host coevolution, and knowledge of the ways in which deployed resistance genes are overcome can guide future resistance development and deployment strategies.

For this reason, we aimed to characterize an Indian *Xoo* strain, IX-221, associated with an outbreak on *xa13*-containing rice in an experimental field in the state of Haryana (Yugander et al., 2017), by fully sequencing the genome to evaluate TALE content and to functionally characterize its major TALEs. We report here that Tal7/PthXo2B_IX-221_ is a major virulence factor in both *japonica* and *indica* rice, yet activates SWEET13 only in japonica rice. It activates no clade III SWEET gene in *indica*. Further, a designer TALE with the same RVD sequence as PthXo2B_IX-221_ acted as a virulence factor only in *japonica* rice, pointing toward an influence of repeat backbone sequence variation on DNA targeting capacity.

## Materials and Methods

### Genome sequencing, assembly and annotation

For sequencing, genomic DNA was extracted and a 20 kb library prepared as previously described (Booher et al., 2015, Carpenter *et al*., 2020). Two single molecule real time (SMRT) cells were used to sequence the library on a PacBio RSII machine (Pacific Biosciences, Menlo Park, CA USA). The sequence reads were *de novo* assembled using HGAP 3.0 and SMRTAnalysis 2.3, and the assembly was verified using local *tal* gene assembly with PBX (Booher et al., 2015) and the variant finder PBHoney from PB Suite 14.7.14 (English *et al*., 2014). The verified whole genome sequence was annotated using the National Centre for Biotechnology Information (NCBI) with Prokaryotic Genome Annotation Pipeline (PGAP) (Tatusova *et al*., 2016).

### Strains, primers, plant material and inoculations

The bacterial strains used in this study were *E coli* DH5), *Xanthomonas oryzae* pv *oryzae* (*Xoo*) strains IX-221, PXO99A (ME2) and *Agrobacterium tumefaciens* (At) GV3101. *E coli* and *At* cells were grown in Luria-Bertani (LB) medium at 37°C and 28°C respectively, *Xoo* strains at 28°C in GYE (20g/l Glucose, 10g/l yeast extract). Plasmids were introduced into *E coli* by heat shock and by electroporation into *Xoo* and *At*. Antibiotics were used at the following concentrations: Ampicillin, 100μg/ml; spectinomycin, 50μg/ml; kanamycin, 25μg/ml; tetracycline, 10μg/ml for *E coli* and 2μg/ml for *Xoo*. Primers used in the study are provided in Table S4.

Rice plants were grown in a growth chamber maintained at 28°C and 85% relative humidity (RH) with a photoperiod of 12h. *Oryza sativa* ssp. *japonica* cv Nipponbare and the near-isogenic *O sativa* ssp. *indica* line IR24 were used for disease assays and gene expression assays as described (Carpenter et al., 2020). *Nicotiana benthamiana* (*Nb*) plants were grown under 16 h of light, 50% RH, at 25:20°C, day: night in the growth chamber. Fully expanded eaves of 5 weeks old plants were inoculated with *At* strains with constructs using a needleless syringe. Significant differences were determined using the paired Student’s t-test. Experiments were repeated thrice.

### TALE analysis, target prediction, cloning and generation of TALE and dTALE constructs

All *tal* gene sequences were extracted and their orthology to previously sequenced TALEs were determined using the AnnoTALE (Richter *et al*., 2014a). The *tal* gene repertoires were verified by Southern blots of genomic DNA digested with *Sph*I, and probed with the *tal* gene specific probe pZWavrXa7.

Targets of Tal2b, Tal6b and Tal7 were predicted using the TALE-NT 2.0 Target Finder tool (Doyle *et al*., 2012). Predictions were made for both forward and reverse strands of promoter sequences, defined 1000 bp upstream of the translational start site for TALE-NT 2.0, and using MSU Rice Genome Annotation Project Release 7 (http://rice.plantbiology.msu.edu/). Default settings were used for Target Finder (upstream base of binding site = T, score cutoff = 3.0, Doyle et al. scoring matrix).

For cloning *tal* genes, a subgenomic library of IX-221 digested with *Bam*HI was generated in pBS II (KS-) and screened for *tal* positive clones by PCR and Sanger sequencing. The Tal2b, Tal6b and Tal7 were further subcloned as *Sph*I fragment into an entry vector pCS466 on the *Sph*I site flanked by N- and C-termini of *Xo* pv oryzicola (*Xoc*) strain BLS256 *tal*1c gene (Verdier et al 2012). The dTALE derivatives of Tal6b and Tal7 were assembled using Golden Gate kit (Cermak *et al*., 2011), into another entry vector pTAL1, also encoding the *Xoc tal*1c without its central repeat region. The complete *tal* and *dtal* genes thus reconstituted were then transferred to broad host range destination vector pKEB31(Cermak et al., 2011) by Gateway LR clonase reaction (Invitrogen/Thermo Fisher Scientific) for expression in *Xoo* strain ME2 and in pGWB5 for expression in *At* strain GV3101.

### GUS assay

GUS reporter constructs were generated by cloning the SWEET13 EBEs of IR24 and Nipponbare on the unique *Asc*I site of binary vector pCS752, having the pepper Bs3 promoter driving *uid*A reporter gene expression with the *Asc*I site upstream of the native AvrBs3 binding element. *At* GV3101 transformed wit TALE and dTALE constructs and GUS reporter constructs were resuspended to an OD600 of 0.8 in 10 mM MgCl_2_ with 150 µM of acetosyringone and mixed 1:1 for inoculation into fully expanded leaves of 5week old *Nicotiana benthamiana* plants. Leaf discs were samples at 48hpi for qualitative and quantitative assays as previously described (Römer *et al*., 2009, Carter *et al*., 2020).

## Results

### IX-221 harbors three major TALEs, each with one or two aberrant repeats

We first confirmed the *xa13* compatibility of IX-221 by inoculating to rice line IRBB13, homozygous for *xa13*, with the near isogenic parent IR24 used as a susceptible control, and found IX-221 indeed to be compatible with *xa13* (data not shown). We then sequenced the whole genome of IX-221 and assembled the data *de-novo*. The assembly yielded a genome consisting of a single, 4.9 Mb circular chromosomal contig with 63.7% G+C content with 172X average sequence coverage. The genome, like those of other *Xoo* strains, contains hundreds of IS elements, which contribute to genomic plasticity (Table S1). The number and sizes of *TALE* genes in the IX-221 genome was confirmed by Southern blot (Figure S1). There are 20 encoded TALEs, of which 18 fall into existing TALE classes as defined by AnnoTALE version. 1.5 (Grau *et al*., 2016) (Table S2), including a TruncTALE (also called iTALE), a class of TALEs with truncated N- and C-termini that suppress resistance mediated by *Xo1* or *Xa1* (Ji *et al*., 2016, Read *et al*., 2016). Of the two IX-221 TALEs classified as new by AnnoTALE, one contains only five RVDs. The other is orthologous with the major TALE PthXo3, based on analysis using FuncTALE, which groups TALEs by predicted DNA target sequences (Pérez-Quintero *et al*., 2015). Strikingly, in addition to the PthXo3 ortholog, IX-221 also harbors two orthologs of another major TALE, PthXo2, and each of these three TALEs has one or two aberrant repeats (Table S2). None of the IX-221 TALEs is a PthXo1 ortholog, consistent with the compatibility of the strain with *xa13*.

The PthXo3 ortholog, Tal2b_IX-221_ (AnnoTALE class IU), like PthXo3, has a 39-aa repeat that in PthXo3 is important for frameshift binding to the target (Richter et al., 2014a). Because it is a variant of PthXo3, distinct from the three observed to date [Oliva, 2019], we hereafter refer to it as PthXo3D_IX-221_. With the full RVD sequence of PthXo3D_IX-221_ as input, using TALE-NT 2.0 (Doyle et al., 2012) and the rice cv. Nipponbare reference genome sequence, no binding site in the *SWEET14* promoter was predicted, but with the RVD of the 39-aa repeat excluded, PthXo3D_IX-221_ was predicted to bind the PthXo3 EBE (Figure S2). As expected, IX-221, and the *pthXo1* mutant derivative ME2 of *Xoo* strain PXO99A (Yang & White, 2004) carrying PthXo3D_IX-221_ on a plasmid, each induced *SWEET14* in Nipponbare leaves (Figure S3). Aligned to the Nipponbare EBE, PthXo3 has a better predicted binding score ratio (Doyle et al., 2012) than PthXo3_IX-221_ (Figure S2). However, PthXo3D_IX-221_ includes more RVDs with relaxed or semi-relaxed base specificity. Namely, it has three NS, an RVD that accommodates A, C, or G; (Yang *et al*., 2014), and one NN, which recognizes G or A; that flexibility may allow activation of yet uncharacterized *SWEET14* alleles with variations at the corresponding positions in the EBE. The flexibility may in fact have been selected for, by the presence of such alleles.

The PthXo2 orthologs are Tal6b_IX-221_ and Tal7_IX-221_. PthXo2 variants reported to date include PthXo2B and PthXo2C (Oliva *et al*., 2019a). While PthXo2 has standard, 34 aa repeats throughout, both PthXo2B and 2C have 36 aa in their 9^th^ and 12^th^ repeats and differ in a few RVDs from PthXo2. Tal6b_IX-221_, with 36 aa only in its 12^th^ repeat is a novel variant. Alignment of the RVD sequences positions Tal6b between PthXo2, and PthXo2B and 2C together (Figure S4). We hereafter refer to Tal6b_IX-221_ as PthXo2D_IX-221_. Tal7_IX-221_ has 36-aa in its 9^th^ and 12^th^ repeats. In fact, it is identical to PthXo2B from strain PXO61 (PthXo2B_PXO61_) except that, relative to Tal7_IX-221_, PthXo2B_PXO61_ has a 1 aa insertion (glycine) 32 aa from the N-terminus (at position 33), in the region associated with type III secretion (Szurek *et al*., 2002), and a 2 aa substitution at positions 31 and 32 of repeat 19. The 2 aa substitution replaces glutamine and aspartic acid with arginine and alanine; while aspartic acid and alanine are each common at position 32 in TALE repeats, the arginine at position 31 is unusual. We hereafter refer to Tal7_IX-221_ as PthXo2B_IX-221_. In all three of the 36-aa repeats in the IX-221 TALEs, the canonical proline and valine residues at positions 29 and 30 are repeated (as a pair).

### PthXo2D_IX-221_ activates SWEET13 and confers virulence on *indica* rice only

Given the seeming redundance of PthXo2D_IX-221_ and PthXo2B_IX-221_ with the *SWEET14* activator PthXo3_IX-221_ in IX-221, we questioned whether these PthXo2 orthologs are in fact activators of *SWEET13.* Beginning with PthXo2D_IX-221_, we first determined the binding score ratio for its RVD sequence on the *SWEET13* allele present in the *japonica* variety Nipponbare and the allele in the *indica* variety IR24, using the target finder tool of TALE-NT 2.0 (Doyle et al., 2012). Because the 36 aa repeat is an aberrant type capable of disengaging (Becker et al., 2022), we determined also the score ratio using the sequence with the RVD of that repeat excluded. Using a score ratio of 3 or less as a cutoff for predicted binding, PthXo2D_IX-221_ is expected to bind well to the IR24 allele with its 36 aa repeat engaged and marginally to the Nipponbare allele with it excluded (PthXo2D and PthXo2D.1, respectively, Figure 1a).

**FIGURE 1.**
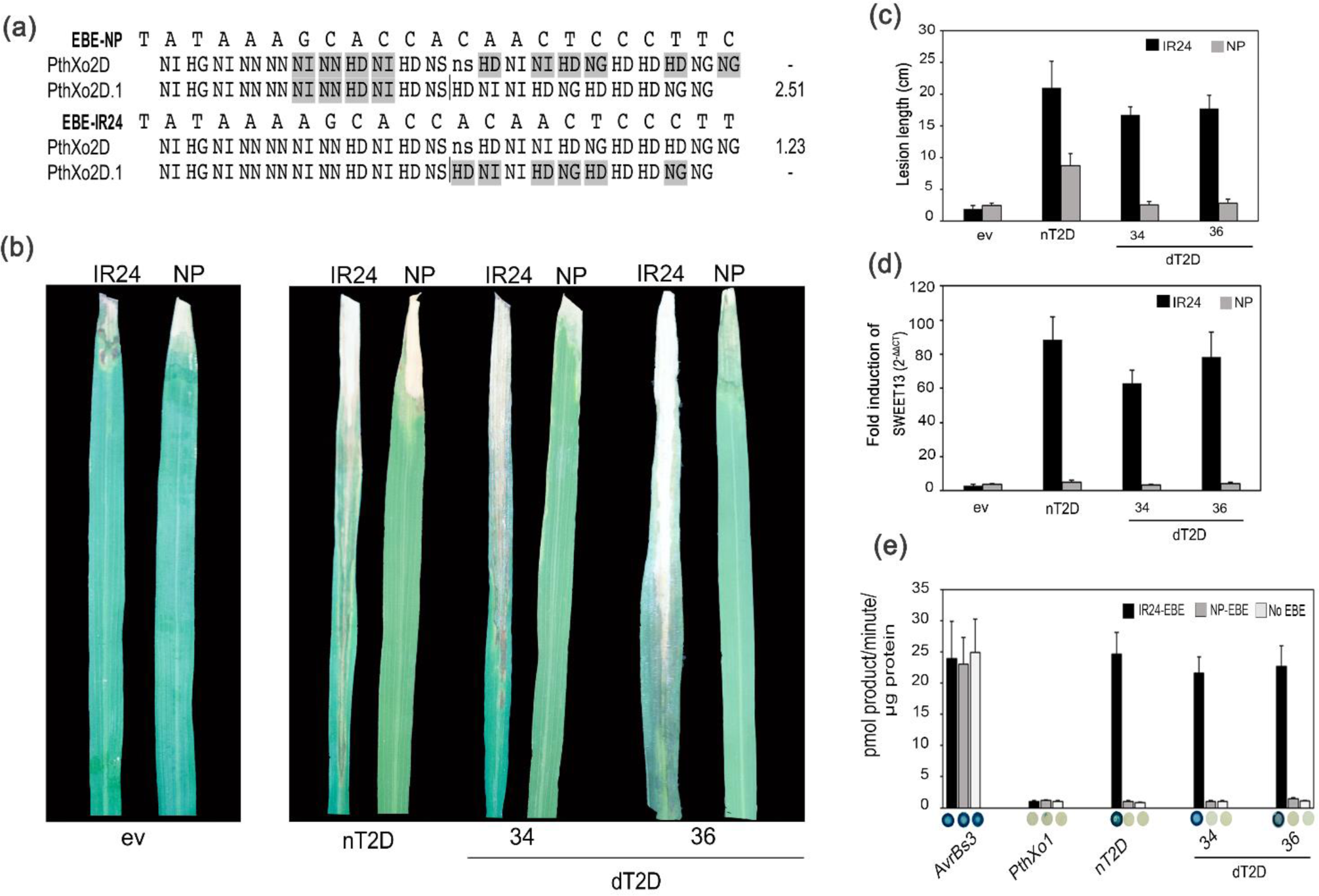
PthXo2D_IX-221_ acts as a major TALE in *indica* rice variety IR24 without relying on its 36-aa aberrant repeat to disengage. (a) TALE-NT 2.0-based prediction of binding of PthXo2D_IX-221,_ and a variant with the aberrant repeat removed, to *SWEET13* alleles found in rice cv. Nipponbare (NP) and IR24. RVDs mismatching the aligned base are highlighted in grey and that of the 36-aa aberrant repeat is lowercase. The position from which the RVD of the aberrant repeat was omitted for the prediction is indicated by a vertical line. A score ratio (ratio of observed EBE score to the best possible score for the TALE) of 3 was used as cut off for predicted binding. A ‘-’ indicates a score above cut-off, i.e., not predicted to bind. (b) Representative images and (c) lesion lengths on leaves of 6-week-old rice (cv. NP and IR24) plants 14 days after clip inoculation with ME2 expressing the indicated TALE or a negative control TALE lacking a central repeat region (pAC99). (d) Expression of *SWEET*13, measured by RT-quantitative real time PCR, in NP and IR24 leaves 24-27 hr following syringe infiltration of ME2 expressing the indicated TALE or control, relative to mock-inoculum. (e) Reporter assay of EBE binding by nT2D and dTALE variants as indicated. *N. benthamiana* leaves were co-infiltrated with *Agrobacterium tumefaciens* strains delivering the indicated TALE or dTALE construct and a GUS reporter construct driven by a minimal *Bs*3 promoter containing the indicated EBE, and GUS activity assayed 48 hr later. Shown are means for three independent infiltrations. Error bars represent standard deviation. A representative leaf disc is shown for each.

Next, to test the predictions, we cloned the native *pthXo2D_IX-221_* central repeat region (as an *SphI* fragment), between the flanking N- and C-terminal regions of Tal1c from *X oryzae* pv. oryzicola strain BLS256 in the expression vector pKEB31 (Cermak et al., 2011), and assembled a designer TALE (dTALE) construct encoding PthXo2D_IX-221_ with its aberrant repeat replaced by a standard repeat, also using the Tal1c context. We designated these constructs nT2D (nT for native TALE) and dT2D_34 (dT for dTALE), respectively. To control for any effect of differences in the dTALE repeat backbone sequences from the native ones, we also generated a dTALE equivalent of PthXo2D_IX-221_ i.e., with its 12^th^ repeat 36 aa in length, dT2D_36. We tested each of these constructs in ME2 inoculated to IR24 and Nipponbare plants. ME2 carrying nT2D and its equivalent dTALE: dT2D_36 induced *SWEET13* and caused long lesions when inoculated to IR24 (Figure 1b, c and d). In Nipponbare, neither detectably induced *SWEET13* or caused long lesions typical of a fully virulent strain. However, ME2 with nT2D did elicit lesions longer than the negative control, ME2 transformed with pAC99, a plasmid encoding a TALE with the CRR removed (Cernadas *et al*., 2014) (Figure 1b and c). d2D_34, with the standard repeat, behaved the same as PthXo2, inducing SWEET13 and restoring virulence to ME2 only in IR24, with no virulence increase relative to the control in Nipponbare. We therefore infer that PthXo2D_IX-221_ is functionally distinct from PthXo2, which has no aberrant repeats and restores virulence to ME2 only in *indica* rice (Zhou *et al*., 2015).

As a complementary approach and to confirm binding, we used *Agrobacterium*-mediated transient transformation in *Nicotiana benthamiana* leaves, as described (Römer et al., 2009), to test whether nT2D, dT2D_34, and dT2D_36 could activate GUS reporter constructs driven by a minimal promoter from the pepper *Bs3* gene (Römer et al., 2009) amended either with the PthXo2 EBE from the Nipponbare allele of *SWEET13* or with the EBE from the IR24 allele. The TALE AvrBs3, which activates the minimal *Bs3* promoter, was used as a positive control, and PthXo1, which has no EBE in either reporter construct, was used as a negative control. nT2D, dT2D_34, and dT2D_36 strongly induced the reporter harboring the IR24 EBE, and not the Nipponbare EBE (Figure 1e), validating the observations made in rice leaves, and also indicating PthXo2D_IX-221_ binds the *SWEET13* allele in IR24 without its 36-aa repeat disengaging.

Why n2D partially rescued ME2 in Nipponbare while d2D_36 did not is unclear. Perhaps n2D, and not d2D_36, activates *SWEET13* marginally enough so as not to be statistically significant but sufficiently to partially restore virulence. Such a difference in activity between the two proteins might derive from minor differences in target affinity due to differences between the native and the designer repeat backbone sequences.

### PthXo2B_IX-221_ renders both *indica* and *japonica* rice susceptible, and does not rely on any Clade III SWEET gene in *indica*

Following the same approach for PthXo2B_IX-221,_ we determined using TALE-NT 2.0 that it is likely to bind only the *japonica* allele, and only when the RVD of one or the other of its 36 aa repeats is excluded (PthXo2B.1 and PthXo2B.2, respectively, Figure 2a). Similar to PthXo2D_IX-221_, the expression constructs assembled for PthXo2B_IX-221_ included the native CRR, a designer equivalent with 36 aa 9^th^ and 12^th^ repeats, and CRRs with one the other or both aberrant repeats converted to standard, 34 aa repeats, nT2B, dT2B_36_36, dT2B_34_36, dT2B_36_34, and dT2B_34_34, respectively.

**FIGURE 2.**
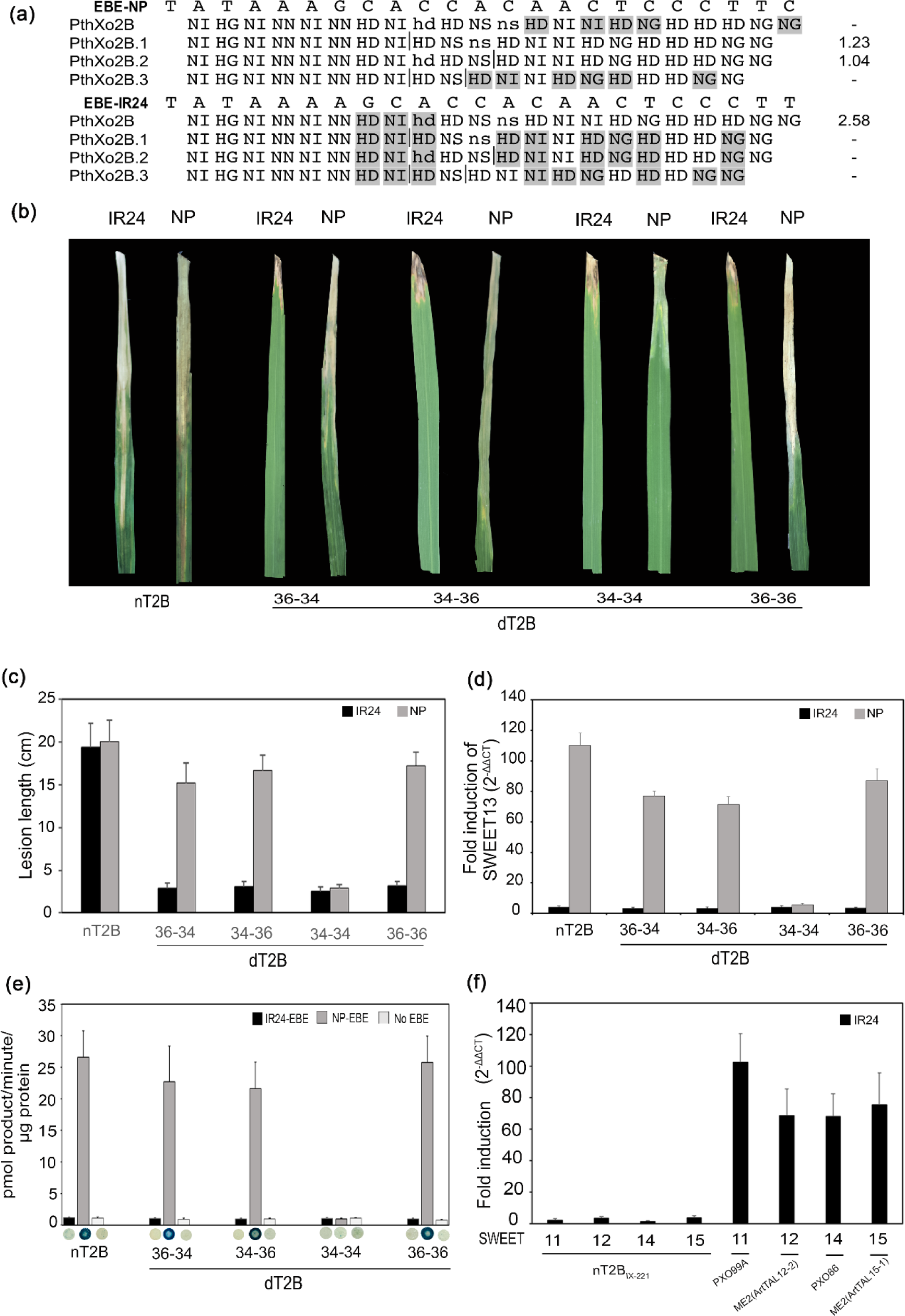
PthXo2B_IX-221_ acts as a major TALE in both *indica* rice variety IR24 and *japonica* variety Nipponbare (NP) relying on either of its 36-aa repeats to disengage in NP and without activating any Clade III SWEET in IR24. TALE-NT 2.0-based prediction of binding of PthXo2B_IX-221_ and variants with one or both aberrant repeats removed, to *SWEET13* alleles of rice cv. Nipponbare (NP) and IR24, as in Figure 1. (b-e) as in Figure 1 with nT2B and dTALE variants as indicated. (f) Fold induction, as in (b), of the other clade III *SWEET* genes by nT2B and selected positive control strains that induce cognate *SWEET* genes.

nT2B in ME2, as predicted, induced *SWEET13* and restored virulence in Nipponbare, and did not upregulate *SWEET13* in IR24 (Figure 2b, c and d). Results with the dTALEs in ME2 indicated that activation of *SWEET13* in Nipponbare depends on either one or the other of the 36 aa repeats disengaging: dT2B_36_36, dT2B_34_36, and dT2B36_34 each activated *SWEET13* and restored virulence to ME2 in Nipponbare, while the dTALE with both repeats replaced, dT2B_34_34, did not. Also, like nT2B, none of the dTALEs activated *SWEET13* in IR24 (Figure 2d). The EBE binding assay results aligned with these observations, with all but dTB_34_34 strongly inducing the GUS reporter with the Nipponbare EBE only, thus confirming that PthXo2B_IX-221_ binds only the Nipponbare allele and that it relies on either one of its two aberrant repeats disengaging to do so (Figure 2e). This conclusion is consistent with that of Becker and colleagues (2022) based on GUS reporter assays using synthetic TalBK2 (AnnoTALE class for PthXo2B) and variants missing either repeat.

Surprisingly, despite nT2B not activating *SWEET13* in IR24, it nonetheless fully restored virulence of ME2 in that variety (Figure 2b and c), and it did so without activating any other clade III *SWEET* gene (Figure 2f). This contrasts with what Xu et al (2019) and Oliva and colleagues (2019a) reported with PthXo2B_PXO61_, which has the same sequence of RVDs as PthXo2B_IX-221_: that TALE did not restore virulence to ME2 in IR24. As noted above, there is a 1 aa insertion in the N-terminal region and a 2 aa substitution in repeat 19 in PthXo2B_PXO61_ relative to PthXo2B_IX-221._ In addition, there are a few minor substitutions in the N-terminal region of PthXo2B_PXO61_ relative to the fragment of Tal1c used for the PthXo2B_IX-221_ expression construct (and the dTALEs). Those substitutions reside in a region that, while essential for type III secretion (Szurek et al., 2002), is dispensable for DNA binding (Miller *et al*., 2011). How PthXo2B_IX-221_ restores virulence to ME2 without activating *SWEET13* is unclear. It does not have a predicted binding site in any other clade III *SWEET* promoter, in either orientation, suggesting involvement of a non-*SWEET* susceptibility target in *indica* varieties. Further exploration, beyond the scope of this study, will be necessary to test that hypothesis. In this context however, we tentatively conclude that the non-canonical substitution at repeat 19 changes the specificity or affinity contribution of that repeat, altering the targeting profile such that PthXo2B_PXO61_ does not activate any alternative S gene.

More surprising still, the PthXo2B_IX-221_-equivalent dTALE dT2B_36_36 failed to restore virulence to ME2 in IR24, just as dT2D_36, the dTALE equivalent of PthXo2D_IX-221_, failed to restore any virulence in Nipponbare, while nT2D, with the native CRR, partially did so. Alignment of the CRRs (Figure S5) revealed some differences. The standard repeat consensus subsequence VAIAS present in the dTALEs is replaced by MAIAN in the native CRRs, in repeats harboring the RVD ‘NN’ (repeats 4 and 6 of PthXo2B_IX-221_ and 4, 5 and 7 of PthXo2D_IX-221_). And, in the 6^th^ and 7^th^ repeats of PthXo2B_IX-221_ and PthXo2D_IX-221_, respectively, the canonical D or A at position 4 is replaced by T. To explore the prevalence of these substitutions, we scanned TALE repeat sequences from diverse randomly picked *Xoo* genomes as well as genomes of the closely related *X. oryzae* pv. oryzicola (Xoc), which causes bacterial leaf streak of rice. We found that the NN repeats of all PthXo2 orthologs, from diverse Xoo strains, have the MAIAN subsequence substitution for VAIAS, and that no other repeats in those orthologs or any repeats in other TALEs do (Table S3). Similarly, we found T at position 4 in all PthXo2 orthologs, and in no other TALEs. Further, we found each of the two subsequences in TALEs of *Xoc* (Table S3), and always in an NN repeat, but never together in the same repeat. We hypothesize that the ‘MAIAN’ and ‘T’ substitutions relative to the dTALEs, in PthXo2B_IX-221_ are important for its ability to act as major TALE in IR24 and in PthXo2D_IX-221_ its ability of to confer some virulence in Nipponbare, ostensibly by altering the targeting profiles to include one or more S genes other than a clade III *SWEET*. Whether this is indeed the case and how, remains to be explored.

## Discussion

The Indian *Xoo* strain IX-221 is armed with three, distinct TALEs that each could allow the strain to cause disease in a host genotype with resistance governed by *xa13*. Importantly, each of the three TALEs has one or more functional aberrant repeats, which could allow them to accommodate *SWEET* allelic variation or to target alternative S genes, or both. The first, PthXo3_IX-221_, is a variant of and functions the same as PthXo3, activating SWEET14 in both *japonica* and *indica*, with its sole aberrant repeat disengaged. It harbours more RVDs with relaxed specificity than PthXo3 however, suggesting that it may be adapted to as yet unidentified *SWEET14* allelic variation, perhaps in landraces not yet characterized. The second is a PthXo2 ortholog, PthXo2B_IX-221_. We have shown that TALE PthXo2B_IX-221_ is capable of activating SWEET13 in *japonica*, and ostensibly an alternative *S* gene in *indica*, to render genotypes in both subspecies susceptible. Strains harbouring this TALE are thereby expected to circumvent the broad spectrum resistance of recently developed rice varieties with edits at known EBEs in the promoters of *SWEET11*, -*13* and -*14* (Xu et al., 2019, Oliva et al., 2019a). Existence of an additional target that functions as an alternative S gene has also been postulated for the *SWEET14* inducer TalC from the African *Xoo* strain BAI3 (Blanvillain-Baufume *et al*., 2017); TalC confers virulence even in a line in which its EBE in *SWEET14* had been disrupted. Unfortunately, because the dTALE equivalent of PthXo2B_IX-221_ did not render ME2 virulent in the *indica* cultivar IR24, we were unable to test with our standard repeat derivatives of that construct whether the aberrant repeats are important for the ability of PthXo2B_IX-221_ to do so. The third TALE in IX-221 that confers an ability to overcome *xa13* is PthXo2D_IX-221_. It functions as predicted, activating the *SWEET13* allele and conferring virulence in the *indica* cultivar IR24. Although the 36-aa repeat of PthXo2D_IX-221_ apparently does not disengage for recognition of the Nipponbare *SWEET13* allele, its presence hints at alleles in landraces, yet unidentified, for which the capacity of this repeat to disengage is important.

The ability of PthXo2B_IX-221_ to function as a major TALE in an *indica* rice genotype, strikingly differentiates it from the previously characterized PthXo2B_PXO61_. We speculate that PthXo2B_PXO61_ does not bind and activate the presumed PthXo2B_IX-221_ target in *indica* sufficiently to render the plant susceptible. The amino acid alignment of the two, points to the Q to R substitution in PthXo2B_PXO61_ as the cause for this. As noted, it may change the affinity contribution of the 19^th^ repeat, reducing affinity overall, or possibly the base specificity of the repeat and thus the targeting profile of the TALE. An alternative, not mutually exclusive possibility is that the substitution makes the 19th repeat like an aberrant one but disengaging in an obligate rather than facultative fashion. Finally, it is formally possible that the difference is due to a lower amount of PthXo2B_PXO61_ in the plant cell since that study used the low copy plasmid pHM1 for PthXo2B_PXO61_ while we used the moderate copy number plasmid pKEB31. However, based on the fact that the repeat backbone differences between the native TALE constructs and their dTALE equivalents in our study also impacted function, we posit that repeat backbone polymorphisms indeed can affect the function of individual repeats and thus the overall targeting profile of a TALE effector. Altogether, the presence of three major TALEs in IX-221, each with one or more repeat types that can facultatively disengage, and one with more lax DNA targeting specificity than its previously characterized ortholog, provide compelling evidence of intense selection pressure on the *Xoo* population, particularly in India, to acquire or evolve TALEs that equip a strain to overcome *xa13*-mediated resistance against bacterial blight disease. Of note, while PthXo2B from each of four additional Philippine strains (Oliva et al, 2019) has the same Q to R substitution in repeat 19 that PthXo2B_PXO61_ does; we found a PthXo2B allele in the Taiwanese strain XM9 that matches PthXo2B_IX-221_, suggesting that the ability to activate the putative alternative S gene is not restricted to Indian strains.

## ACKNOWLEDGEMENTS

This work was supported by Department of Biotechnology award BT/CEIB/12/1/01 and the Indian Council of Agricultural Research-National Project on Functional Genomics and Genetic modification in crops to RR. RR thanks Rohini Sreevathsa, M Rathinam and Narsimha Dokka, ICAR-NIPB, New Delhi, India for help and support.

## DATA AVAILABILITY STATEMENT

Genome data for IX-221 generated in this study are available through Genbank Accession number CP019228.

## REFERENCES

Antony, G., Zhou, J., Huang, S., Li, T., Liu, B., White, F., et al. (2010) Rice xa13 recessive resistance to bacterial blight is defeated by induction of the disease susceptibility gene Os-11N3. The Plant cell, tpc. 110.078964.

Becker, S., Mücke, S., Grau, J. and Boch, J. (2022) Flexible TALEs for an expanded use in gene activation, virulence and scaffold engineering. Nucleic Acids Res., 50, 2387–2400.

Blanvillain-Baufume, S., Reschke, M., Sole, M., Auguy, F., Doucoure, H., Szurek, B., et al. (2017) Targeted promoter editing for rice resistance to *Xanthomonas oryzae* pv. oryzae reveals differential activities for SWEET14-inducing TAL effectors. Plant Biotechnol J, 15, 306–317.

Boch, J., Scholze, H., Schornack, S., Landgraf, A., Hahn, S., Kay, S., et al. (2009) Breaking the code of DNA binding specificity of TAL-type III effectors. Science, 326, 1509–1512.

Bogdanove, A. J., Schornack, S. and Lahaye, T. (2010) TAL effectors: finding plant genes for disease and defense. Current opinion in plant biology, 13, 394–401.

Booher, N. J., Carpenter, S. C., Sebra, R. P., Wang, L., Salzberg, S. L., Leach, J. E., et al. (2015) Single molecule real-time sequencing of Xanthomonas oryzae genomes reveals a dynamic structure and complex TAL (transcription activator-like) effector gene relationships. Microbial genomics, 1.

Carpenter, S. C., Mishra, P., Ghoshal, C., Dash, P. K., Wang, L., Midha, S., et al. (2020) An xa5 resistance gene-breaking Indian strain of the rice bacterial blight pathogen Xanthomonas oryzae pv. oryzae is nearly identical to a Thai strain. Frontiers in microbiology, 11, 579504.

Carter, M. E., Carpenter, S. C., Dubrow, Z. E., Sabol, M. R., Rinaldi, F. C., Lastovetsky, O. A., et al. (2020) A TAL effector-like protein of an endofungal bacterium increases the stress tolerance and alters the transcriptome of the host. Proceedings of the National Academy of Sciences, 117, 17122–17129.

Cermak, T., Doyle, E. L., Christian, M., Wang, L., Zhang, Y., Schmidt, C., et al. (2011) Efficient design and assembly of custom TALEN and other TAL effector-based constructs for DNA targeting. Nucleic Acids Res, 39, e82.

Cernadas, R. A., Doyle, E. L., Nino-Liu, D. O., Wilkins, K. E., Bancroft, T., Wang, L., et al. (2014) Code-assisted discovery of TAL effector targets in bacterial leaf streak of rice reveals contrast with bacterial blight and a novel susceptibility gene. PLoS pathogens, 10, e1003972.

Chu, Z., Fu, B., Yang, H., Xu, C., Li, Z., Sanchez, A., et al. (2006) Targeting xa13, a recessive gene for bacterial blight resistance in rice. TAG. Theoretical and applied genetics. Theoretische und angewandte Genetik, 112, 455–461.

Denancé, N., Szurek, B., Doyle, E. L., Lauber, E., Fontaine-Bodin, L., Carrère, S., et al. (2018) Two ancestral genes shaped the Xanthomonas campestris TAL effector gene repertoire. New Phytol., 219, 391–407.

Doyle, E. L., Booher, N. J., Standage, D. S., Voytas, D. F., Brendel, V. P., VanDyk, J. K., et al. (2012) TAL Effector-Nucleotide Targeter (TALE-NT) 2.0: tools for TAL effector design and target prediction. Nucleic acids research, 40, W117–W122.

English, A. C., Salerno, W. J. and Reid, J. G. (2014) PBHoney: identifying genomic variants via long-read discordance and interrupted mapping. BMC bioinformatics, 15, 180.

Grau, J., Reschke, M., Erkes, A., Streubel, J., Morgan, R. D., Wilson, G. G., et al. (2016) AnnoTALE: bioinformatics tools for identification, annotation, and nomenclature of TALEs from *Xanthomonas* genomic sequences. Sci Rep, 6, 21077.

Hutin, M., Pérez-Quintero, A. L., Lopez, C. and Szurek, B. (2015) MorTAL Kombat: the story of defense against TAL effectors through loss-of-susceptibility. Frontiers in plant science, 6, 535.

Ji, Z., Ji, C., Liu, B., Zou, L., Chen, G. and Yang, B. (2016) Interfering TAL effectors of *Xanthomonas oryzae* neutralize R-gene-mediated plant disease resistance. Nat Commun, 7, 13435.

Lore, J. S., Vikal, Y., Hunjan, M. S., Goel, R. K., Bharaj, T. S. and Raina, G. L. (2011) Genotypic and Pathotypic Diversity of *Xanthomonas oryzae* pv. oryzae, the Cause of Bacterial Blight of Rice in Punjab State of India. Journal of Phytopathology, 159, 479–487.

Midha, S., Bansal, K., Kumar, S., Girija, A. M., Mishra, D., Brahma, K., et al. (2017) Population genomic insights into variation and evolution of *Xanthomonas oryzae* pv. oryzae. Sci Rep, 7, 40694.

Miller, J. C., Tan, S., Qiao, G., Barlow, K. A., Wang, J., Xia, D. F., et al. (2011) A TALE nuclease architecture for efficient genome editing. Nature biotechnology, 29, 143–148.

Mishra, D., Vishnupriya, M. R., Anil, M. G., Konda, K., Raj, Y. and Sonti, R. V. (2013) Pathotype and genetic diversity amongst Indian isolates of *Xanthomonas oryzae* pv. oryzae. PLoS One, 8, e81996.

Mondal, K. K., Meena, B. R., Junaid, A., Verma, G., Mani, C., Majumder, D., et al. (2014) Pathotyping and genetic screening of type III effectors in Indian strains of Xanthomonas oryzae pv. oryzae causing bacterial leaf blight of rice. Physiol. Mol. Plant Pathol., 86, 98–106.

Moscou, M. J. and Bogdanove, A. J. (2009) A simple cipher governs DNA recognition by TAL effectors. Science, 326, 1501–1501.

Nino-Liu, D. O., Ronald, P. C. and Bogdanove, A. J. (2006) Xanthomonas oryzae pathovars: model pathogens of a model crop. Molecular plant pathology, 7, 303–324.

Oliva, R., Ji, C., Atienza-Grande, G., Huguet-Tapia, J. C., Perez-Quintero, A., Li, T., et al. (2019a) Broad-spectrum resistance to bacterial blight in rice using genome editing. Nature biotechnology, 37, 1344–1350.

Oliva, R., Ji, C., Atienza-Grande, G., Huguet-Tapia, J. C., Perez-Quintero, A., Li, T., et al. (2019b) Broad-spectrum resistance to bacterial blight in rice using genome editing. Nat. Biotechnol., 37, 1344–1350.

Pérez-Quintero, A. L., Lamy, L., Gordon, J. L., Escalon, A., Cunnac, S., Szurek, B., et al. (2015) QueTAL: a suite of tools to classify and compare TAL effectors functionally and phylogenetically. Front. Plant Sci., 6, 545.

Read, A. C., Rinaldi, F. C., Hutin, M., He, Y. Q., Triplett, L. R. and Bogdanove, A. J. (2016) Suppression of Xo1-Mediated Disease Resistance in Rice by a Truncated, Non-DNA-Binding TAL Effector of *Xanthomonas oryzae*. Front Plant Sci, 7, 1516.

Richter, A., Streubel, J., Blucher, C., Szurek, B., Reschke, M., Grau, J., et al. (2014a) A TAL effector repeat architecture for frameshift binding. Nat Commun, 5, 3447.

Richter, A., Streubel, J., Blücher, C., Szurek, B., Reschke, M., Grau, J., et al. (2014b) A TAL effector repeat architecture for frameshift binding. Nature communications, 5, 1–10.

Römer, P., Recht, S. and Lahaye, T. (2009) A single plant resistance gene promoter engineered to recognize multiple TAL effectors from disparate pathogens. Proceedings of the National Academy of Sciences, 106, 20526–20531.

Streubel, J., Pesce, C., Hutin, M., Koebnik, R., Boch, J. and Szurek, B. (2013) Five phylogenetically close rice SWEET genes confer TAL effector-mediated susceptibility to *Xanthomonas oryzae* pv. oryzae. New Phytol, 200, 808–819.

Szurek, B., Rossier, O., Hause, G. and Bonas, U. (2002) Type III-dependent translocation of the Xanthomonas AvrBs3 protein into the plant cell. Molecular microbiology, 46, 13–23.

Tatusova, T., DiCuccio, M., Badretdin, A., Chetvernin, V., Nawrocki, E. P., Zaslavsky, L., et al. (2016) NCBI prokaryotic genome annotation pipeline. Nucleic acids research, 44, 6614–6624.

Wilkins, K. E., Booher, N. J., Wang, L. and Bogdanove, A. J. (2015) TAL effectors and activation of predicted host targets distinguish Asian from African strains of the rice pathogen Xanthomonas oryzae pv. oryzicola while strict conservation suggests universal importance of five TAL effectors. Front. Plant Sci., 6, 536.

Xu, Z., Xu, X., Gong, Q., Li, Z., Li, Y., Wang, S., et al. (2019) Engineering broad-spectrum bacterial blight resistance by simultaneously disrupting variable TALE-binding elements of multiple susceptibility genes in rice. Molecular plant, 12, 1434–1446.

Yang, B., Sugio, A. and White, F. F. (2006) Os8N3 is a host disease-susceptibility gene for bacterial blight of rice. Proceedings of the National Academy of Sciences of the United States of America, 103, 10503–10508.

Yang, B. and White, F. F. (2004) Diverse members of the AvrBs3/PthA family of type III effectors are major virulence determinants in bacterial blight disease of rice. Molecular plant-microbe interactions, 17, 1192–1200.

Yang, J., Zhang, Y., Yuan, P., Zhou, Y., Cai, C., Ren, Q., et al. (2014) Complete decoding of TAL effectors for DNA recognition. Cell Res., 24, 628–631.

Yu, Y., Streubel, J., Balzergue, S., Champion, A., Boch, J., Koebnik, R., et al. (2011) Colonization of rice leaf blades by an African strain of Xanthomonas oryzae pv. oryzae depends on a new TAL effector that induces the rice nodulin-3 Os11N3 gene. Molecular Plant-Microbe Interactions, 24, 1102–1113.

Yugander, A., Sundaram, R. M., Ladhalakshmi, D., Hajira, S. K., Prakasam, V., Prasad, M. S., et al. (2017) Virulence profiling of *Xanthomonas oryzae* pv. oryzae isolates, causing bacterial blight of rice in India. European Journal of Plant Pathology, 149, 171–191.

Zhou, J., Peng, Z., Long, J., Sosso, D., Liu, B., Eom, J. S., et al. (2015) Gene targeting by the TAL effector PthXo2 reveals cryptic resistance gene for bacterial blight of rice. The Plant journal: for cell and molecular biology, 82, 632–643.

