## Supplemental Information for "A PthXo2B ortholog in *Xanthomonas oryzae* pv oryzae strain IX-221 acts as a major virulence factor on *indica* rice without activating a Clade III *SWEET* gene"

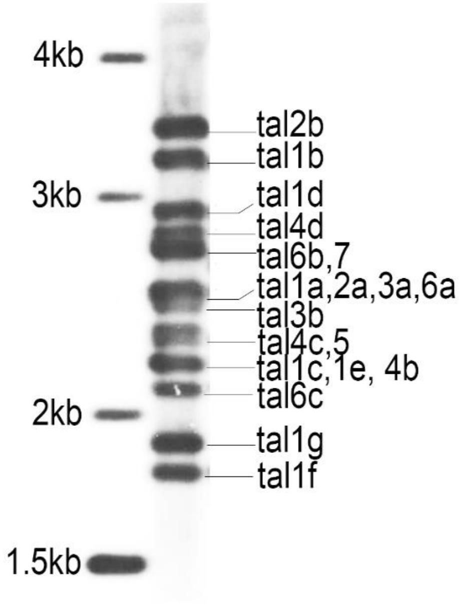


**FIGURE S1. Southern blot showing *TALE* gene content in IX-221.** *Sph*I-digested genomic DNA of IX-221 was probed with *avrXa7.*

| | A | T | A | T | A | A | A | C | C | C | C | C | T | C | C | A | A | C | C | A | G | G | T | G | C | T | A | A | G | Score ratio |
|---|---|---|---|---|---|---|---|---|---|---|---|---|---|---|---|---|---|---|---|---|---|---|---|---|---|---|---|---|---|---|
| PthXo3D$_{IX-221}$ | NI | HG | NI | HG | NI | NI | NI | HD | NN | HD | HD | HD | NS | NI | n* | NI | NS | HD | NS | NS | NN | NN | NN | HG | NN | HD | N* | NS | NG | 2.5 |
| PthXo3 | NI | HG | NI | HG | NI | NI | NI | HD | NN | HD | HD | HD | NG | HD | n* | NI | HD | HD | NN | NS | NI | NN | NN | NG | NN | HD | N* | NS | N* | 2.2 |

**FIGURE S2**. **RVD sequences of PthXo3 and PthXo3D$_{IX-221}$ aligned with the EBE in *SWEET14*.** RVDs in blue differ from corresponding RVDs in PthXo3$_{PXO61}$. The RVD of the 39-aa repeat is lowercase. An asterisk indicates that the second amino acid in the RVD is absent, resulting in a 33aa repeat. RVDs highlighted in grey are mismatched with the base in the EBE. Score ratio is the ratio of the actual Target Finder score to the ideal score. Score ratios shown are for RVD sequences excluding the 39-aa repeat; with the repeat included, both scores were above the Target Finder score ratio cutoff of 3.

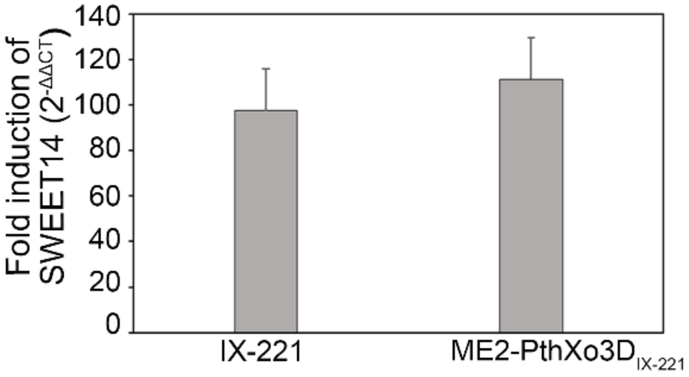


**FIGURE S3**. **PthXo3D$_{IX-221}$ activates *OsSWEET14*.** Expression of *OsSWEET14* in rice cv. Nipponbare leaves measured by RT-qPCR in response to inoculation by syringe infiltration with IX-221 and ME2 expressing PthXo3D$_{IX-221}$, relative to mock-inoculated leaves. Leaf tissues were sampled 24-27 hrs after inoculation.

| | 1 | 2 | 3 | 4 | 5 | 6 | 7 | 8 | 9 | 10 | 11 | 12 | 13 | 14 | 15 | 16 | 17 | 18 | 19 | 20 | 21 | 22 |
|---|---|---|---|---|---|---|---|---|---|---|---|---|---|---|---|---|---|---|---|---|---|---|
| PthXo2 | NI | HG | NI | NN | NN | NI | NN | HD | NI | HD | NS | NS | NS | HD | NN | HD | NG | HD | HD | HD | NG | NG |
| PthXo2D | NI | HG | NI | NN | NN | NI | NN | HD | NI | HD | NS | ns | HD | NI | NI | HD | NG | HD | HD | HD | NG | NG |
| PthXo2C | NI | HG | NI | NN | NI | NN | HD | HD | hd | HD | NS | ns | HD | NI | NI | HD | NG | HD | HD | HD | NG | NG |
| PthXo2B | NI | HG | NI | NN | NI | NN | HD | NI | hd | HD | NS | ns | HD | NI | NI | HD | NG | HD | HD | HD | NG | NG |

**FIGURE S4**. **PthXo2D$_{IX-221}$ is a novel PthXo2 ortholog more similar to PthXo2 than described PthXo2 variants.** An alignment of RVD sequences is shown. RVDs in blue differ from RVDs at the corresponding positions in in PthXo2. RVDs of 36-aa repeats are lowercase.

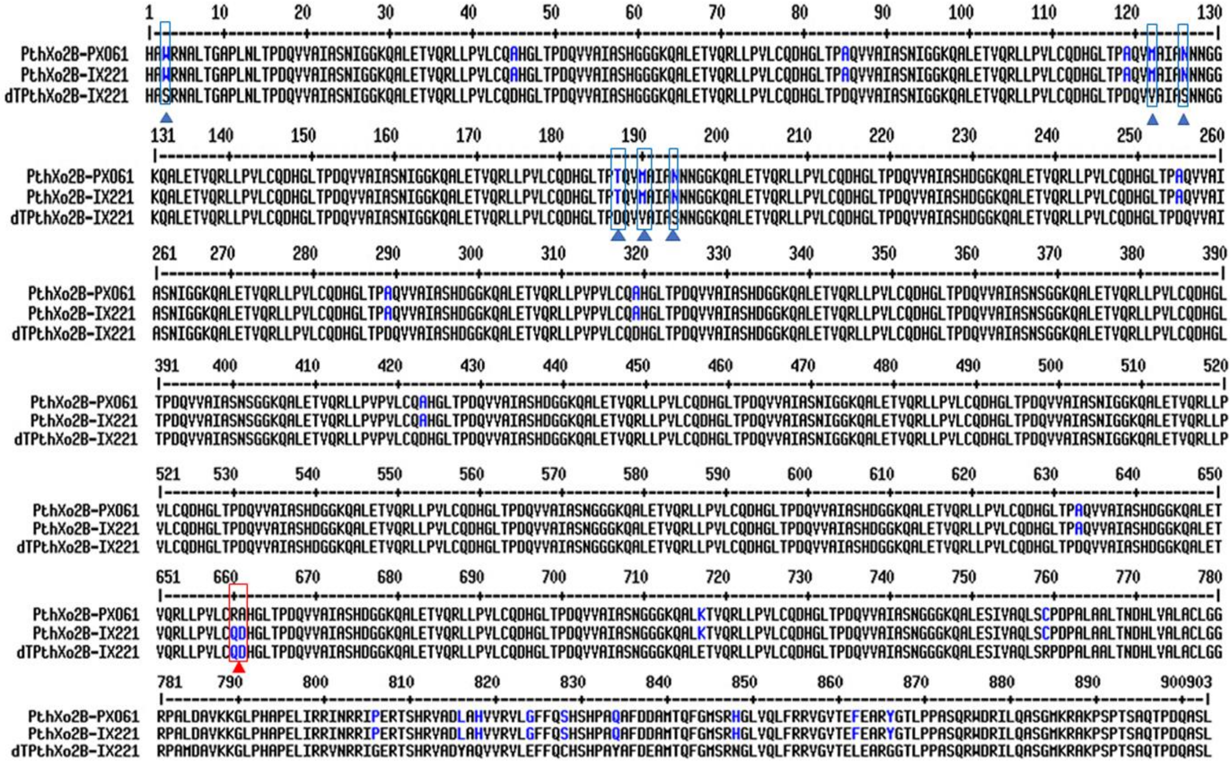

**FIGURE S5**. **Sequence alignment of PthXo2B$_{IX\text{-}221}$ with its ortholog and dTALE equivalent highlights non canonical polymorphisms in the repeat backbone.** Alignment of the amino acid sequences in the central repeat regions (delimited by *Sph*I sites) of PthXo2B$_{IX\text{-}221}$, PthXo2B$_{PXO61}$ and dTALEPthXo2B$_{IX\text{-}221}$ 36-36 is shown. Non-canonical amino acid substitutions in the native PthXo2 proteins relative to the standard amino acids in the dTALE are in blue font. The single non canonical amino acid in the 19$^{th}$ repeat of PthXo2B$_{PXO61}$ differentiating it from PthXo2B$_{IX\text{-}221}$ and its dTALE is in red font.

**TABLE S1**. **Genome assembly/ characteristics**

| | **IX-221** |
|---|---|
| *Genome* | |
| Length (bp) | 4,944,063 |
| GC content (%) | 63.7 |
| Annotated genes | 4,987 |
| IS elements (complete/fragments) | 416/102 |
| *Assembly* | |
| Mapped N50 (bp) | 14818 |
| Average coverage depth | 172x |

### TABLE S2. RVD sequences and AnnoTALE classes of TALEs encoded in the IX-221 genome

| Name | RVD Sequence[a] | | | | | | | | | | | | | | | | | | | | | | | | | | | | | Class | Ortholog |
|---|---|---|---|---|---|---|---|---|---|---|---|---|---|---|---|---|---|---|---|---|---|---|---|---|---|---|---|---|---|---|---|
| | 1 | 2 | 3 | 4 | 5 | 6 | 7 | 8 | 9 | 10 | 11 | 12 | 13 | 14 | 15 | 16 | 17 | 18 | 19 | 20 | 21 | 22 | 23 | 24 | 25 | 26 | 27 | 28 | 29 | | |
| Tal1g | NI | HG | NS | NN | NN | HD | NI | HD | NN | HG | NS | N* | HD | NG | | | | | | | | | | | | | | | | BJ | AvrXa10 |
| Tal1f | NI | NN | NI | HG | HG | HD | NG | HD | HG | HD | HD | HD | NG | | | | | | | | | | | | | | | | | AE | |
| Tal1e | NI | NN | N* | NG | NS | NN | NN | NN | NI | NN | NI | NG | HD | HD | NI | HG | N* | | | | | | | | | | | | | AO | AvrXa27 |
| Tal1d | NN | HD | NS | NG | HD | NN | N* | NI | HD | NS | HD | NN | HD | NN | HD | NN | NN | NN | NN | NN | NN | NN | HD | NG | | | | | | AD | |
| Tal1c | NI | NS | HD | NG | NS | NN | HD | H* | NN | NN | HI | NN | HD | NG | HD | HD | N* | | | | | | | | | | | | | AL | |
| Tal1b | HD | HD | NN | NN | NS | NG | HD | S* | HG | HD | NG | N* | HD | HD | HD | N* | NN | nq[1] | NN | HD | HI | ND | HD | HG | NN | HG | N* | | | AQ | AvrXa23 |
| Tal1a | HD | HD | HD | NG | N* | HD | HD | HD | N* | NI | NI | NN | NN | HD | NG | HD | NI | HD | NG | NG | | | | | | | | | | AP | |
| Tal2a | NI | HG | NI | NI | NI | NN | HD | NS | NN | NS | NN | HD | NN | HI | HD | NN | NI | NG | HD | NG | | | | | | | | | | AB | |
| Tal2b | NI | HG | NI | HG | NI | NI | NI | HD | NN | HD | HD | HD | NS | NI | n*[2] | NI | NS | HD | NS | NS | NN | NN | NN | HG | NN | HD | N* | NS | NG | IU | **PthXo3D** |
| Tal3a | NI | N* | NI | NS | NN | NG | NN | NS | N* | NS | NN | NS | N* | NI | HG | HD | NI | HD | HD | NG | | | | | | | | | | AH | |
| Tal3b | NI | HG | NI | HG | HG | HD | NS | NG | HD | NG | NG | HG | HG | HD | HG | HD | HD | NN | NG | | | | | | | | | | | AA | |
| Tal4a | NS | NI | HD | NS | NG | | | | | | | | | | | | | | | | | | | | | | | | | FM | |
| Tal4b | NI | HG | NI | NS | NN | HG | NN | NI | HD | HD | NG | HD | NS | NI | N* | NS | NG | | | | | | | | | | | | | CA | |
| Tal4c | NI | HG | NI | NI | NN | HD | NN | HD | NS | HD | SS | HD | NI | NI | NN | NI | NN | NG | | | | | | | | | | | | IT | |
| Tal4d | NI | H* | NI | NN | NN | NN | NN | NN | HD | NI | NS | HG | HD | NI | N* | NS | NI | NI | HD | HD | N* | NS | N* | | | | | | | AR | PthXo6 |
| Tal5 | NS | NG | NG | ng[3] | NG | NG | HD | HD | HD | HD | NN | HD | NG | HD | HD | HD | NN | H* | | | | | | | | | | | | AI | TruncTALE |
| Tal6a | NI | NG | NN | NG | NK | NG | NS | NN | NI | NN | NI | NN | NS | NG | NS | NN | NI | N* | NS | NG | | | | | | | | | | AG | |
| Tal6b | NI | HG | NI | NN | NN | NI | NN | HD | NI | HD | NS | ns[4] | HD | NI | NI | HD | NG | HD | HD | HD | NG | NG | | | | | | | | AM | **PthXo2D** |
| Tal6c | NI | NS | HD | HG | NS | NN | HD | H* | NG | NN | NN | HD | HD | HD | HD | NG | | | | | | | | | | | | | | BA | |
| Tal7 | NI | HG | NI | NN | NI | NN | HD | NI | hd[4] | HD | NS | ns[4] | HD | NI | NI | HD | NG | HD | HD | HD | NG | NG | | | | | | | | BK | **PthXo2B** |

**TABLE S3. Presence/absence of the repeat subsequence MAIAN and LTPT in TALEs of *Xoo* and *X. oryzae* pv. oryzicola strains.**

| Pathovar | Strain | TALE with MAIAN | TALE with LTPT | RA in repeat 19 of PthXo2B | Accession number |
|---|---|---|---|---|---|
| *Xoo* | BAI3 | none | none | n/a | CP025610.1 |
| *Xoo* | AXO1947 | none | none | n/a | CP013666.1 |
| *Xoo* | MAI1 | none | none | n/a | CP025609.1 |
| *Xoo* | MAI106 | none | none | n/a | CP019089.1 |
| *Xoo* | MAI129 | none | none | n/a | CP019090.1 |
| *Xoo* | MAI134 | none | none | n/a | CP019089.1 |
| *Xoo* | MAI145 | none | none | n/a | CP019092.1 |
| *Xoo* | MAI68 | none | none | n/a | CP019085.1 |
| *Xoo* | MAI73 | none | none | n/a | CP019086.1 |
| *Xoo* | MAI95 | none | none | n/a | CP019087.1 |
| *Xoo* | MAI99 | none | none | n/a | CP019088.1 |
| *Xoo* | IX-280 | none | none | n/a | CP019226.1 |
| *Xoo* | PXO79 | none | none | n/a | CP031462.1 |
| *Xoo* | PXO145 | none | none | n/a | CP013961.1 |
| *Xoo* | PXO211 | none | none | n/a | CP013674.1 |
| *Xoo* | PXO524 | none | none | n/a | CP013677.1 |
| *Xoo* | PXO83 | none | none | n/a | CP012947.1 |
| *Xoo* | PXO86 | none | none | n/a | CP007166.1 |
| *Xoo* | PXO99[A] | none | none | n/a | CP000967.2 |
| *Xoo* | SK2-3 | none | none | n/a | CP019515.1 |
| *Xoo* | CFBP1948 | none | none | n/a | CP033185.1 |
| *Xoo* | CFBP1949 | none | none | n/a | CP033184.1 |
| *Xoo* | CFBP1951 | none | none | n/a | CP033183.1 |
| *Xoo* | CFBP1952 | none | none | n/a | CP033182.1 |
| *Xoo* | CFBP7319 | none | none | n/a | CP033181.1 |
| *Xoo* | CFBP7320 | none | none | n/a | CP033186.1 |
| *Xoo* | CFBP7322 | none | none | n/a | CP033179.1 |
| *Xoo* | CFBP7323 | none | none | n/a | CP033178.1 |
| *Xoo* | CFBP7337 | none | none | n/a | CP033175.1 |
| *Xoo* | CFBP7340 | none | none | n/a | CP033174.1 |
| *Xoo* | CFBP8172 | none | none | n/a | CP033173.1 |
| *Xoo* | CIX2374 | none | none | n/a | CP036377.1 |
| *Xoo* | Dak16 | none | none | n/a | CP033172.1 |

| | | | | | |
|---|---|---|---|---|---|
| *Xoo* | T19 | none | none | n/a | CP033171.1 |
| *Xoo* | NXO260 | none | none | n/a | CP033192.1 |
| *Xoo* | PXO282 | none | none | n/a | CP013676.1 |
| *Xoo* | Ug11 | none | none | n/a | CP033170.1 |
| *Xoo* | PXO602 | PthXo2 | PthXo2 | n/a | CP013679.1 |
| *Xoo* | PXO339 | PthXo2 | PthXo2 | n/a | CP092382.1 |
| *Xoo* | PXO71 | PthXo2 | PthXo2 | n/a | CP013670.1 |
| *Xoo* | ScYc-b | PthXo2 | PthXo2 | n/a | CP018087.1 |
| *Xoo* | XF89b | PthXo2 | PthXo2 | n/a | CP011532.1 |
| *Xoo* | YC11 | PthXo2 | PthXo2 | n/a | CP031464.1 |
| *Xoo* | XM9 | PthXo2B | PthXo2B | No | CP020334.1 |
| *Xoo* | PXO61 | PthXo2B | PthXo2B | Yes | CP033187.3 |
| *Xoo* | PXO513 | PthXo2B | PthXo2B | Yes | CP033188.1 |
| *Xoo* | PXO364 | PthXo2B | PthXo2B | Yes | CP033191.1 |
| *Xoo* | PXO404 | PthXo2B | PthXo2B | Yes | CP033190.1 |
| *Xoo* | PXO421 | PthXo2B | PthXo2B | Yes | CP033189.1 |
| *Xoo* | KACC10331 | PthXo2C | PthXo2C | n/a | AE013598.1 |
| *Xoo* | KXO85 | PthXo2C | PthXo2C | n/a | CP033197.1 |
| *Xoo* | JW11089 | PthXo2C | PthXo2C | n/a | CP033193.2 |
| *Xoc* | CFBP7342 | Tal4b | Tal4a | n/a | CP007221.1 |
| *Xoc* | GX01 | Tal4b | Tal4a | n/a | CP043403.1 |
| *Xoc* | BLS256 | Tal1b | Tal9a | n/a | CP003057.2 |

**TABLE S4. List of primers used and their sequences.**

| | Primer name | Primer sequence | Purpose |
|---|---|---|---|
| 1 | TALE 1 Fwd | GCC AAG TCC TGC CCG CGA | For screening subgenomic library for *tal* genes |
| 2 | TALE 2 Rev | CCT CCA GGG CGC GTG C | For screening subgenomic library for *tal* genes |
| 3 | 18S Fwd | CTACGTCCCTGCCCTTTGTACA | qPCR |
| 4 | 18S Rev | ACACTTCACCGGACCATTCAA | qPCR |
| 5 | SWT 11 3' UTR Fwd | GACGACAGATTCTCGCTACTG | qPCR |

| | | | |
|---|---|---|---|
| 6 | SWT 11 3' UTR Rev | TGACTGACTGACTGACTGACTGAC | qPCR |
| 7 | SWT 12 BY Fwd | TGATGACCACGACTGACCAGAG | qPCR |
| 8 | SWT 12 BY Rev | AGTACACGTGTCCCCAATCATAC | qPCR |
| 9 | SWT 13 BY FWD | CTCCAGGGCAAACTTGGAGGAGAC | qPCR |
| 10 | SWT13 BY REV | GGGCAGTTGTGATTGATTGA | qPCR |
| 11 | SWT 14 HUTIN Fwd | ACTTGCAAGCAAGAACAGTAGT | qPCR |
| 12 | SWT 14 HUTIN Rev | ATGTTGCCTAGGAGACCAAAGG | qPCR |
| 13 | SWT 15 5'UTR Fwd | ACTGACTGCTCAGTGTGTTT | qPCR |
| 14 | SWT 15 5'UTR Rev | CGATCTCCCTAGAACTCTTC | qPCR |
